# A Proposed Unified Interphase Nucleus Chromosome Structure: Preliminary Preponderance of Evidence

**DOI:** 10.1101/2021.10.08.463051

**Authors:** John Sedat, Angus McDonald, Hu Cang, Joseph Lucas, Muthuvel Arigovindan, Zvi Kam, Cornelis Murre, Michael Elbaum

## Abstract

Cellular cryo-electron tomography (CET) of the cell nucleus using Scanning Transmission Electron Microscopy (STEM) and the use of deconvolution (DC) processing technology has highlighted a large-scale, 100–300 nm interphase chromosome structure (LSS), that is present throughout the nucleus. This chromosome structure appears to coil the nucleosome 11-nm fiber into a defined hollow structure, analogous to a Slinky (S) (1, motif used in 2) helical spring. This S architecture can be used to build chromosome territories, extended to polytene chromosome structure, as well as to the structure of Lampbrush chromosomes.

**Significance Statement:** Cryo-preservation of the nuclear interior allows a large scale interphase chromosome structure—present throughout the nucleus—to be seen for the first time. This structure can be proposed to be a defined coiled entity, a Slinky. This structure can be further used to explain polytene chromosome structure, an unknown chromosome architecture as well as for lampbrush chromosomes. In addition, this new structure can be further organized as chromosome territories, using all 46 human interphase chromosomes as an example, easily into a 10 micron diameter nucleus. Thus, interphase chromosomes can be unified into a flexible defined structure.

## Introduction

A recent publication described deconvolution (DC) of STEM tomography of cryo-preserved cellular structures (3 and references therein). In brief, cells were plunge-frozen, preserving the aqueous cellular structures in a glassy ice medium, then micron-sized slabs were selected from thin regions of whole cells, followed by STEM tomography, and DC data processing (3). The DC substantially filled the missing wedges, resulting from incomplete tilts, regions that had in the past greatly compromised the Z resolution, allowing much greater ease in interpretation of the tomograms. One of the examples of this technology was a tomogram from an area of the nucleus. A large scale interphase chromosome structure was highlighted (3).

This paper further documents and analyzes the chromosome structures seen in these nucleus tomograms. This paper is in four parts: The first part presents evidence – preliminary compelling evidence – for a unified interphase chromosome structure. The second part of this paper presents a proposed unified interphase chromosome architecture. A third part shows that this interphase chromosome structure could be further organized as chromosome territories, using the 46 human chromosomes as examples, to easily fit into a 10-micron diameter nucleus. A fourth part unifies this structure into polytene chromosome architecture and lampbrush chromosomes. Finally the paper ends with a living light microscopy cell study showing that a G1 nucleus has very similar structures throughout this organelle.

A case can be made that the primary data for this paper has very different attributes, such as close spaced 3-dimensional pixels with subtle grey level differences and textures, requiring new ways to display its details and architecture. This is going to be a general problem for many areas (CET and MRI data, for example). The issue is that one has preserved the cell, for example, and all the water, proteins, lipids/membranes, organelles, and nuclear structures are confined together in a dense small volume packed tightly. The usual methods to visualize, for example moving up and down in Z, do not adequately show the 3-dimensional relationships. We propose extensive use of stereo, with extensions, to ameliorate this problem. Evidence for the chromosome structure is primarily in 3-D stereo movies of various kinds (Fig. 1 and RASPa–c’; RASP—rocking angular stereo pairs). Stereo movies have visualization problems that are carefully discussed in the Supporting Information.

**Fig. 1-.**
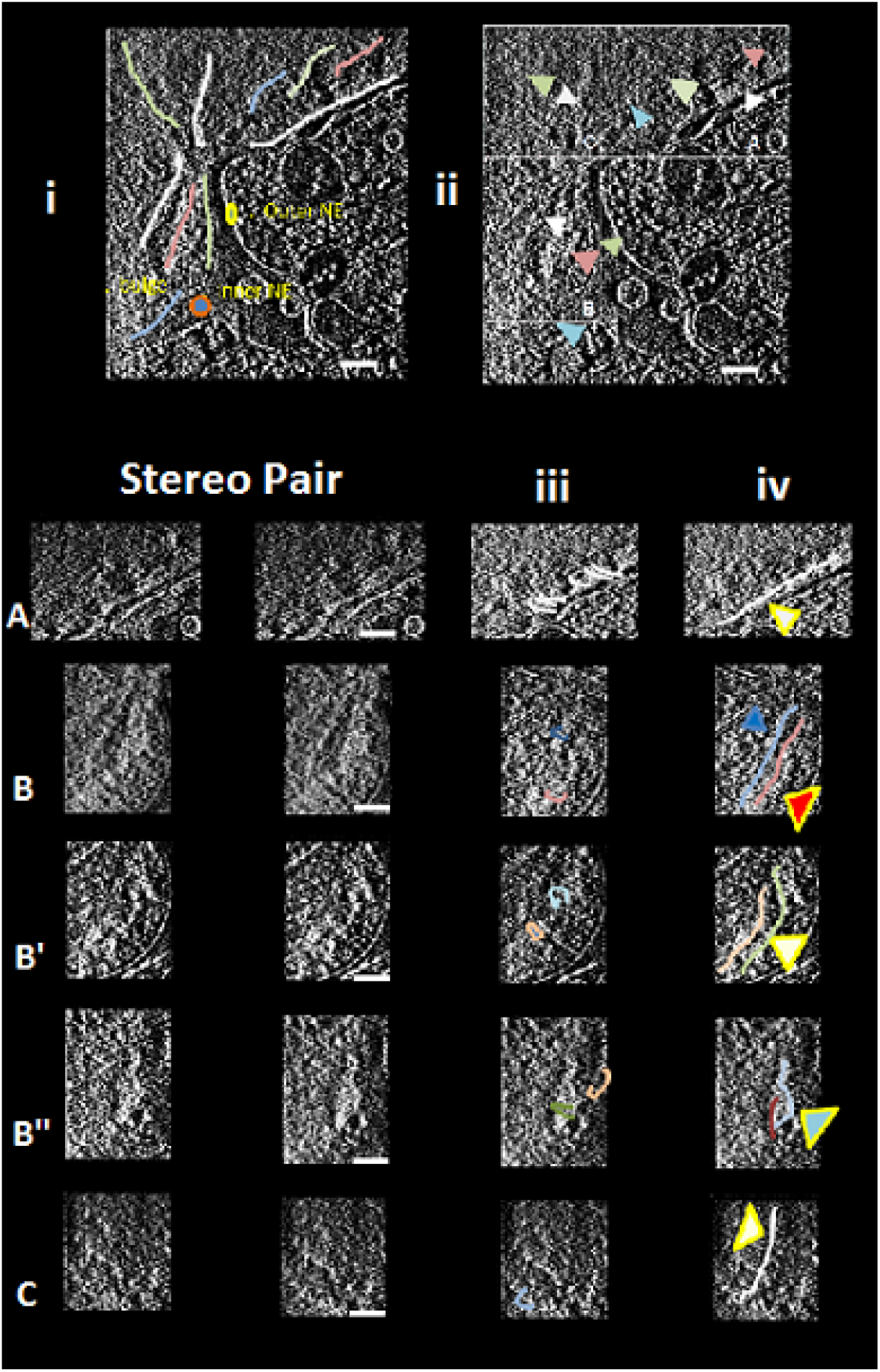
guide. Large scale banded structures are seen in DC of the CET nucleus. Fig. 1 (.mov seen in QuickTime) is 3-dimensional data cube with the ability to move up and down the Z cube axis with the use of the tab. This tab is displaceable to prevent occlusion and coded to record Z position as described in the text.(Fig 1 is located in the Supplement with the RASPs) Fig. 1-*guide* is a guide for visualization of the large scale banded chromosome structures. Fig. 1-*guide* i is a guide to some of the cell biology, while Fig. 1-*guide* ii are structures referenced in Fig. 1A and Table I that are located in the nuclear boxes A–C. The lower part of Fig. 1-*guide*, as stereo images for the boxed nuclear regions, marked A–C at various Z nuclear depths (see Table II); column 2B iv marks LSSs, while 1-guide iii marks banded regions referenced in Table II. Scale bar = 400 nm. Abrev.: Euchr=Euchromatin, int.=internal, x-s= cross-section, cyl.=cylindrical. Drawn band loops, Fig. 1-*guide* iii are shown, and LSS regions are drawn in Fig. 1-*guide* iv.

The interphase nucleus encloses the genomic DNA, as well as machineries for regulation of gene expression, RNA synthesis, and DNA replication. The DNA is packaged into chromatin, whose structure in vivo remains enigmatic. While mitotic chromosomes are highly condensed, interphase chromosomes decondense but remain in distinct territories with little overlap. Interphase chromatin is organized in a number of ways, including immutable gene-rich and gene-poor domains in the primary sequence and expression-promoting or suppressing regions that may vary during the cell cycle or across the spectrum of cell differentiation (10). A classical distinction is drawn between euchromatin and heterochromatin, with the former more “open” and prone to expression and the latter more “closed” and prone to silencing (but see 11). However, there is scant proof that different methods, e.g., fluorescence or electron microscopy, or post-translational histone modifications, actually concur in their identification (11).

The predominant model to describe the path of the DNA strand in the interphase nucleus is that of a constrained random walk (13) or fractal globule (14,15). At prophase the dispersed polymers must recondense without entangling. During mitosis, the space-filling interphase chromatin condenses into a compact banded form. These structures are still undefined as ordered or disordered.

The double-stranded DNA polymer itself, which in isolation appears as a semi-flexible, right-hand helix of 2 nm diameter, winds tightly around core histones to form nucleosomes. Each nucleosome has a DNA footprint of 146 base pairs and a geometric diameter of ≈ 11 nm (17–19). The nucleosomes appear as beads on a string, but the density and spacing of nucleosomes along the DNA sequence may be highly variable (20). At the next stage, the nucleosomes were supposed to coil up into a 30-nm filament, possibly as a tight solenoid or alternatively with a zigzag structure (21, 22). The 30-nm filament should then wind more tightly into a 100-nm filament that winds up in the mitotic chromosome (16). The 30-nm filament is today largely considered an artifact, however (23, 24). It is observed in vitro, in isolated or ruptured nuclei, and in cases of deliberate manipulation of divalent cation concentration, but not in intact nuclei (23– 25).

Current insight into chromatin structure arises primarily from methods based on sequencing. With a number of significant variations, chromatin is cross-linked, cleaved, captured, and sequenced in order to determine which sequences lie in close proximity (14, 28). These methods have revealed a structure of topologically associated domains (TADs) of genes whose regulation is controlled concomitantly even if they appear to be distant in sequence (28, 29). While extensive mathematical modeling can generate a 3D map of the genome overall, it is difficult to assess their fidelity in the intermediate domain. They may suffer from cross-linking artifacts similar to microscopy, and only rarely can the analysis be performed on single cells, so that deduced structures represent an average over the cell population. Their unique strength is, of course, the incorporation of sequence into structure, to which microscopy is largely blind (28, 29).

The native structure of chromatin is not known. Different microscopy methods label primarily the oligonucleotide or associated proteins, and chemical cross-linking, i.e., fixation, may stabilize transient interactions artificially or even lead to collapse of soft structures (25). Fluorescence imaging traditionally suffered from poor resolution. Moreover, DNA intercalating dyes may stretch or stiffen the polymer, while fluorescent histones carry the known risks associated with the bulky fluorescent protein tag. Modern super-resolution imaging methods generally require chemical fixation to accommodate long exposures. Traditional electron microscopy also requires fixation, followed by solvent-based dehydration and impregnation with a hardening polymer. Heavy metal salts are added, and their scattering of electrons generates the image contrast. However, they typically bind more strongly to the protein components of chromatin than to DNA itself, so the interpretation is ambiguous (26). This limitation was circumvented recently by use of a DNA binding dye to generate radical oxygen; this induces a localized polymerization of diamino-benzydine, which in turn binds an osmium stain (24). This multi-stage approach is perhaps unique in identifying the DNA component of the chromatin by electron microscopy, though brings its own set of procedure problems. A modest density difference between euchromatin and heterochromatin was found, but there was no evidence for long-range order (25).

Cryo-microscopy and cryo-tomography have also been applied to chromatin studies. These have been rather difficult to interpret, however, due to complications inherent in the image formation mechanism itself. Contrast is generated by electron-optical interference, and the main scattering elements, the nucleosomes, are not very far removed in scale from the limiting resolution that can be reached without averaging. At the large scale, the conventional defocus phase contrast method used in cryo-EM has a poor contrast transfer for low spatial frequencies (27), i.e., for large structures. Even the difference between mitotic chromosomes and interphase DNA, chromatin for example, is difficult to discern on the basis of image contrast.

Cryo-scanning transmission electron tomography (CSTET) is a relatively new addition to the tool chest of cellular imaging techniques. Its most obvious advantages vis-a-vis conventional defocus phase contrast TEM are the ability to accommodate thicker specimens (1-2 microns) and the quantitative contrast based on electron scattering cross-sections. The convenient incoherent bright-field acquisition mode provides a unipolar optical transfer function with the specimen in focus, a long depth of field, and most importantly, strong contrast for low spatial frequencies (27). We have recently demonstrated the application of CSTET in combination with 3D iterative deconvolution processing to whole cell tomography and obtained a view of the cell nucleus that revealed unexpected large-scale structures (3).

## Results Part I: The Preponderance of Experimental Evidence for a Defined Interphase Nuclear Chromosome Structure

### Deconvolved CET Nuclear Data: First Glimpse

The primary experimental evidence for a new interphase nuclear chromosome structure comes from a STEM CET nuclear data set that was deconvolved (3) and is shown in Fig. 1 (this figure is located in the Supplement with the RASPs), but see Fig. 1-guide. The nucleus CET data set originated from a cultured fibroblast cell line, prepared as described in detail in the recent publication (3). This nucleus shows a bulge/polyp protruding from the nucleus with distortions of the nuclear outer membranes (see Fig. 1b), a known perturbation of the nucleus with nuclear lamin network perturbations(34 and references therein).

Fig. 1 is a 3-dimensional DC nucleus data cube with the ability to rapidly move up and down in nucleus Z plane by the Quick Time bar movement (with a number on the bar that correlates to Z position – see figure legend). The nucleus is divided up into three box areas, A–C, to define nuclear subregions. Additional description of this nucleus is shown in Fig. 1-guide (top panels).

It can be seen that there is in many places, in this nucleus, a large-scale structure (LSS) – a structure with a variable size width of 100–300 nm and many microns in length; the LSS(s) are highlighted in Fig. 1-guide top panels. This structure bends and twists along its length as a thick rope-like structure, and is present throughout the nucleus. Both heterochromatin (next to the NE) and euchromatin (in the interior) of the nucleus seem to be built of the same structures. Study of the LSS structure suggests that it is banded with transverse bands (∼100–300 nm), both for heterochromatin and euchromatin. Table 1 highlights many of these banded LSS structures at specific Z planes, interpreting the LSS positions. If one carefully moves up and down in Z, for each boxed area (A–C), the banded LSS(s), extending for long distances, stand out. These banded LSS(s) are further detailed in stereo pair movies below.

**Table I.**
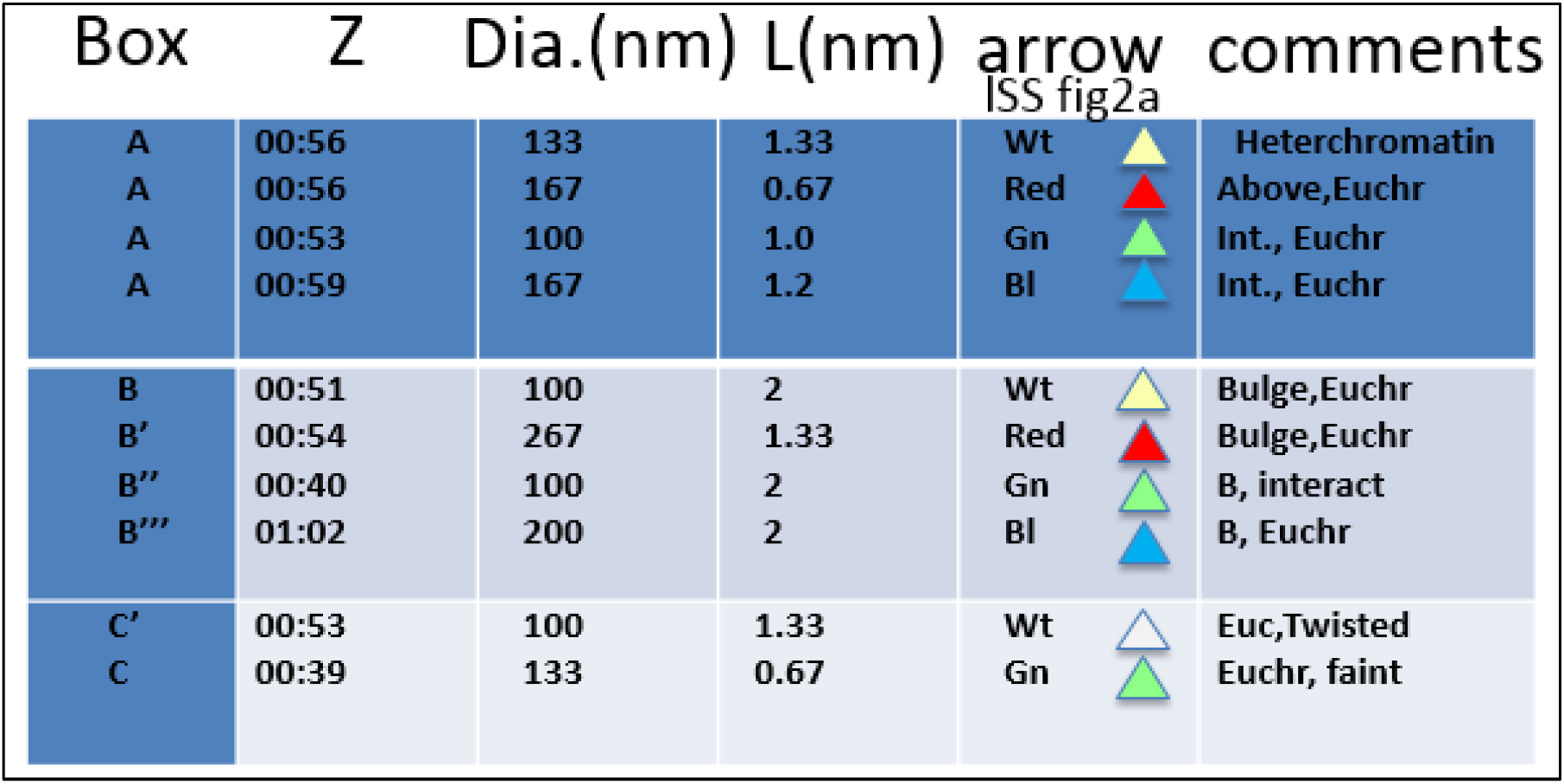
records Z position, location, and diameter dimensions for LSSs shown in Fig2a. This table is studied in conjunction with Fig2b. The LSS are marked by colored arrows. Abrev.: Euchr=Euchromatin, int=internal.

| Box | Z | Dia.(nm) | L(nm) | arrow | comments |
| --- | --- | --- | --- | --- | --- |
| A | 00:56 | 133 | 1.33 | Wt | Heterchromatin |
| A | 00:56 | 167 | 0.67 | Red | Above,Euchr |
| A | 00:53 | 100 | 1.0 | Gn | Int., Euchr |
| A | 00:59 | 167 | 1.2 | Bl | Int., Euchr |
| B | 00:51 | 100 | 2 | Wt | Bulge,Euchr |
| B' | 00:54 | 267 | 1.33 | Red | Bulge,Euchr |
| B'' | 00:40 | 100 | 2 | Gn | B, interact |
| B''' | 01:02 | 200 | 2 | Bl | B, Euchr |
| C' | 00:53 | 100 | 1.33 | Wt | Euc,Twisted |
| C | 00:39 | 133 | 0.67 | Gn | Euchr, faint |

### Detailed Architecture of the LSS Banded Structures

The RASP(s) are now used for the careful analysis of the LSS bands. Fig. 1-guide, and their RASPs corresponding to the nuclear boxes A–C, at a precise Z depth, are detailed. The top two panels in Fig.1-guide show lines for LSS to highlight distinct structures (see figure legend).

A summary of the banded LSSs, from the different nuclear regions, from Fig. 1-guide is shown in Table 2. Fig. 1-guide lower right-hand panels have copies of the stereo pairs, with arrows, so that study of the RASPs can focus on those bands. The reader is advised to study these RASPs carefully, using all the tricks outlined in the Supplement; the data is subtle, but with effort, the structures can be discerned.

**Table II.**
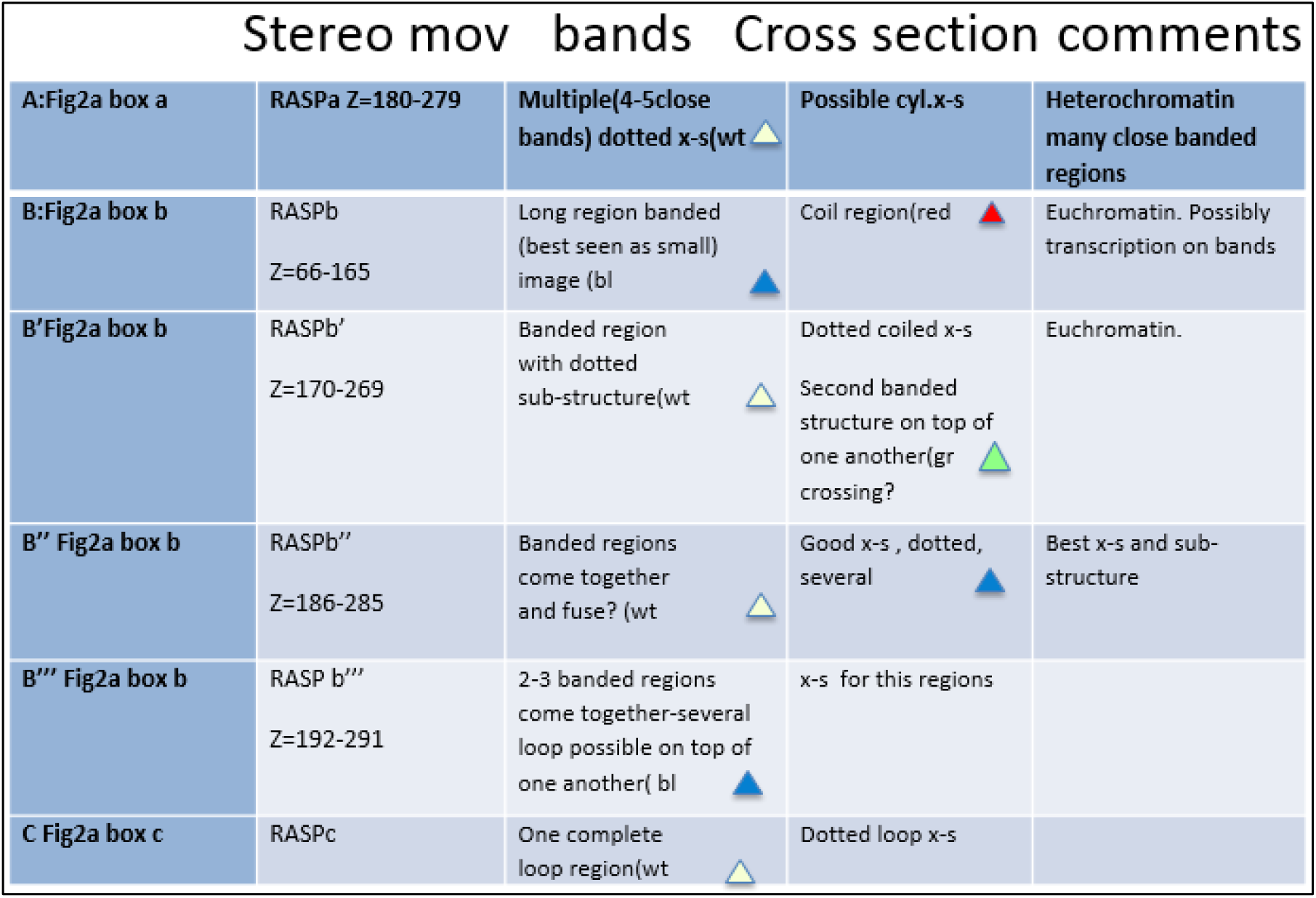
documents RASP stereo movies used to study LSS and their bands--with possible substructure--in the nuclear marked boxes, A-C (see Fig2b). The band locations are labeled with colored arrows. Abrev.: x-s=cross-section, cyl=cylindrical.

Nuclear box A is the first chromosome structure study, a region that contains largely heterochromatin structures, detailed in Fig. 1-guide, and studied in RASPa. The structural features are summarized in Table 2. The heterochromatin LSSs (arrows) show fine punctuated closely packed bands estimated to be 20 nm or less across, inclined (the inclined band size/diameter measures about 300 nm) and appear to be packed in phase with one another. The rims suggest relatively uniform sized dots across many bands for many microns in length. One of the large tilted views (arrows) suggests possible cylindrical cross-sections for the H bands.

Next, the LSSs in the bulge region (largely euchromatin) of the nucleus, Box B (and the different Z depths B–B’’’) are described, and studied by the RASPs. Large number of LSSs, many at different depths, with lengths of many microns come into this bulge. If one walks down these LSSs, the transverse bands stand out. Many have densely packed banded transverse regions (see arrows), many with curving arcs or horse-shoe/cylindrical structures. Some are broken up with thickenings at places in the arcs, possibly coming from transcription events, making it hard to complete the arc. In B’ and B’’, a particularly banded region can be highlighted (arrows), with extensive dotted structures (suggestive of nucleosomes) following a misshapen coiled band(s). Note that many of the arc/horse-shoe band structures are bent/twisted as they seem to coil around. In B’ and B’’, a dense banded region (arrows) is seen, and is likely two or three LSSs on top of each other; study suggests a coiled region in the middle (arrow). Is this region a putative TAD structure (27, 28)? There are regions where it is difficult – so broken up – to really understand the architecture of the LSSs.

A complication for the LSSs and their banded loops is that there are regions, especially in the LSSs and their banded loops, that seem to suggest two structures, very similar to each other in bending and twists, helically wound around each other. We do not know if such possibility could reflect replication, homologue association (see 35) or other complications of chromosome structure.

Fig. 1-guide RASPc (but see especially RASPc’) has highlighted (arrow) an almost complete coil of a dotted chain with a diameter of 200-300nm (depending on how one measures) moving down a suggestive LSS.

A summary stereo pair, Fig. 2, at somewhat higher magnification, shows a representative series of band loops coiled about each other. Dotted coiled band structure (again possible NS) is also highlighted (arrows).

**Figure 2.**
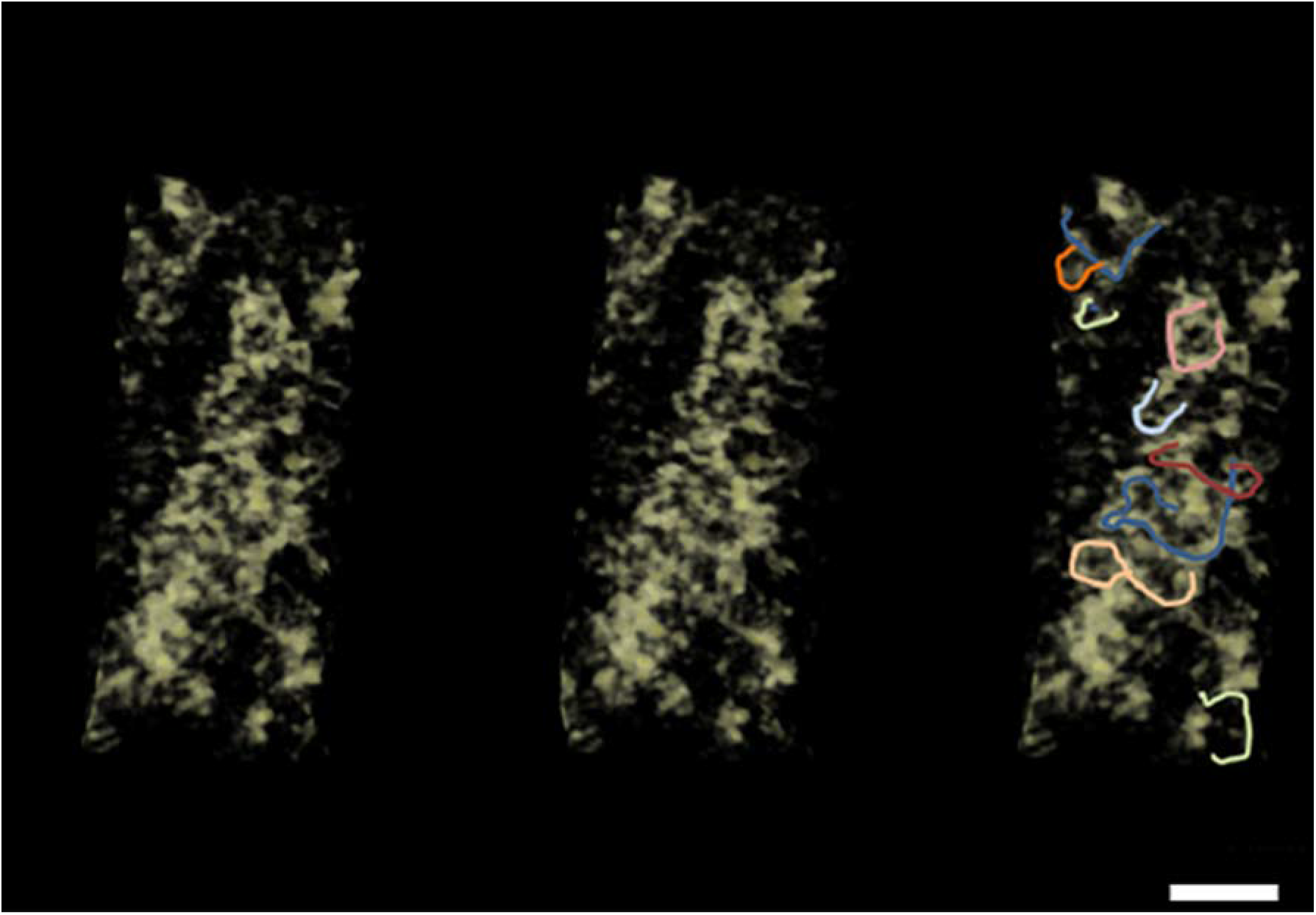
A representative stereo-pair, cut from nuclear box b, of multi-banded chromosome region shows distorted looped (possibly) coiled structures (with substructure). Scale bar=∼200nm. Drawn band loops are shown in the 3^rd^ panel. This figure used UCSF Chimera software to make the stereo image (61).

In summary, the entire interphase nucleus is built of closely packed LSSs – with diameter of 100–300 nm – and transversely banded (likely coiled) structures, a rather regular defined architecture.

## Results Part II: The Proposed Interphase Chromosome Structure

We now integrate the LSS and their transverse bands into a molecular structure. We will use a package of software, described in the Methods, to model, in a quantitative way, multiple sequential levels of helical coiling as structure. The dimensions are accurately tracked and scaled at all levels of the structure. Once the structures are built, they can be analyzed in any orientation, size, or dimension. Table III, a quantitative perspective describes the proposal. First, human diploid DNA, based on sequence, has a total length of 2.02 m (Table IIIA). A representative average human chromosome (chromosome 10) would have a DNA length of 46 mm; as it is bare 2-nm DNA, its compaction is defined as 1 (Table 3B). Essentially all DNA, on average, is organized as nucleosomes (200 base pairs/NS), and chromosome 10 NS chain, if fully extended, would have a shortened length of 19.7mm (see to Fig3 & Table 3 and legend). This further compaction – organized as a well-known 11-nm NS fiber – would have about 43% the DNA length (see Fig 3A & Table 3C). The NS fiber can be fully extended (extended linkers),or a compressed NS fiber with looped linker sequences as a hairpin (Fig. 3A &legend). We propose that this NS 11-nm fiber is further tightly coiled as a 100–300 nm diameter structure (average diameter = 200 nm) whose diameter is the LSS and the transverse bands the coils of the NSs (Table IIID). This tightly coiled structure is analogous to a toy known as a Slinky (1, motif introduced in 2) (defined henceforth as S). S has many attributes and represents the structure we propose for the interphase chromosome architecture. This structure further compacts the DNA by 1/131to 1/393^th^ the length (depending on how tight the NS are packed into the S bands and diameter (S average diameter =200nm with compaction ∼1/260^th^) variations giving rise to a S structure length, for chromosome 10, of 351microns to 117 microns (see Fig3 & Table 3 &legend). Prophase and mitotic chromosome structure is proposed using the S architecture in a subsequent paper.

**Table 3.**
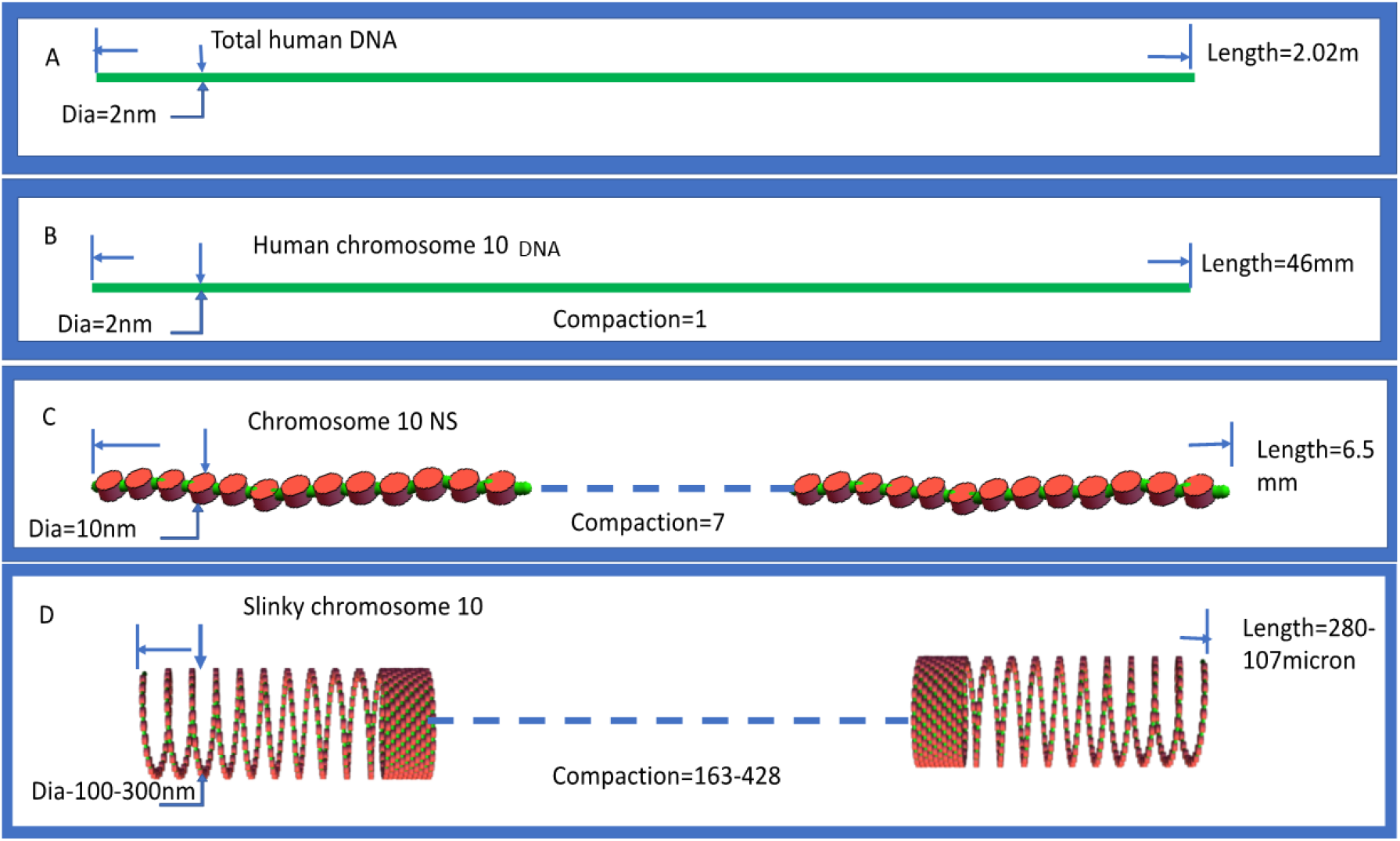
is a depiction of different degrees of organization of DNA in the interphase nucleus. Total human DNA is shown in A. Human chromosome 10, as DNA, --a representative chromosome--is shown in B. Human chromosome 10 is organized as a 10nm NS fiber, as shown in C.(In the literature, the NS diameter is 10nm, but the actual crystal diameter is 11nm(56). The maximal extension of the NS fiber is [669K NS x (11nm NS + 18.4nm(the extended linker DNA))], while the compact NS fiber is [669K NS X (11nm(the NS) +(2×2nm (the DNA) for hairpin bent linker))]. The NS fiber for human chromosome 10 is then coiled as a S with representative diameter of 200 nm, as seen in D. The compaction is S length / DNA length, where the S length is:[(10mm,the compressed NS fiber length)/ S gyrus circumference(to give the number of S gyri)] X 11nm(the gyrus size assuming maximal S packing].

**Figure 3.**
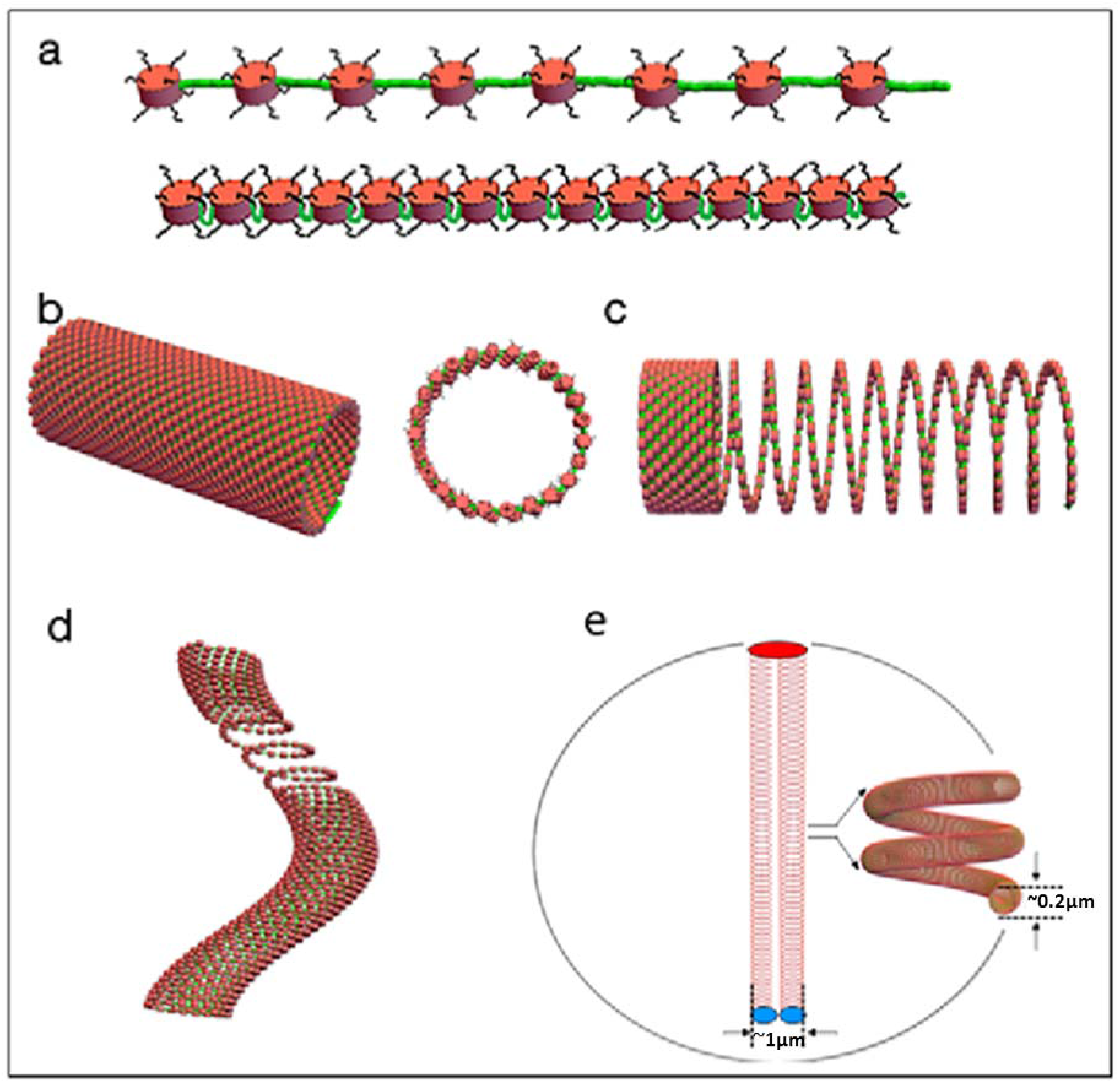
The Slinky as a chromosomal structure motif to explain LSS and its bands. Fig3a shows NSs, to scale, with green linker DNA maximally extending the NS. Underneath shows NS compressed with the linkers looped out, like a hairpin; the looped linkers could act as a molecular antenna with special sequences facing out of the hairpin. Fig3b show the NS coiled (in this case with a diameter of 100nm) as a S. The middle S structure shows a hollow cross-section (dia=100nm), while at a S diameter of 200nm and with maximally condensed NSs would have 63 NS/S turn. The third structure of Fig3c shows S compressed as possible for Heterochromatin or pulled out as for transcribed regions. A bent S structure with a pulled-out active transcribed region is shown in Fig3d. Fig 3e shows human chromosome 10, a metacentric chromosome, coiled as a S, but organized in a 10micron dia. nucleus (to scale), requiring extensive bending and twisting of the S structure(show in the nucleus insert) to fit in the nucleus in the Rable (36,37) configuration.

Diagrammatically, the S structure is depicted in Fig. 3. Part A is the NS 11-nm fiber, to scale, with the NS linker regions extended. Alternatively, these linker regions can be looped out as molecular antennas, in which case the 11-nm NS fiber is more compressed. Part B shows how the NS fiber is coiled as an S, and can be quite regularly packed. Each gyrus could carry between 24 NS and 63 NS, depending on the gyrus diameter and the NS density. The corresponding lengths of DNA are 4.8 kb to 12.6 kb, corresponding to proteins encoding 1600– 4200 amino acids. If the S gyri are closely packed, this could represent heterochromatin. The interior of a cross-section is hollow, allowing for essentially the interior and exterior to be largely topologically equivalent for free diffusion of transcription machinery and/or DNA replication events. Note that the histone tails can be extended equivalently spatially to the interior and exterior. Fig. 3C shows how the S can be pulled out for transcription, a little or a lot, depending on the degree of transcription. Indeed an S gyrus or two can be completely extended out as a looped structure. Pulled out S structures can be bounded by more closely/densely packed S regions. The beauty of the S structure is that it is very flexible and can be distorted with ease such as squashing the S to flatten, slanting the gyri for packing and bending, or bending/distortions of the looped out S regions during transcription etc events.

## Results Part III: Global Nuclear Architecture; Chromosome Territories

The S structure has to fit into a 5–10+ micron nucleus. Fig. 3D shows chromosome 10, to scale, fitting in a 10-micron nucleus. Chromosome 10, an average sized human chromosome, is a metacentric chromosome: the middle centromere region divides the chromosome into two arms. Many interphase chromosomes are organized so that the centromere is located on the nuclear envelope on one end of the nucleus, while the telomere end is at the opposite nucleus pole, the well-known Rabl configuration (36, 37). The S length(s) for chromosome 10 (Table IIID), arranging in the Rabl configuration, indicate that some local bending/twisting/folding of the S structure has to be made to accommodate this length in the nucleus, as shown in Fig. 3e. In a real nucleus this further coiling is likely to be irregularly bending and twisting, though still in a Rabl configuration. One gives this chromosome about one to two microns for this twisting as 45 other chromosomes, possible in S structures, will also have to be accommodated. Nevertheless, the genesis of a chromosome territory (38) is evident. Does the S structure fit comfortably in a 10-micron diameter average sized nucleus? If the 2 m of human genome DNA is maximally coiled into a 200-nm diameter S (1/266the DNA length), it would take up 25% or so of the nuclear volume. This leaves room for S gyri expansion for transcription etc. This fraction of the S volume relative to nuclear volume goes rapidly down as the nuclear diameter increases. Nevertheless, the DNA in a S structure could be somewhat closely packed, and this is what one observes in live imaging microscopy, see Fig. 5.

Do all 46 human interphase chromosomes, in a S structure, – compressed as shown in Fig. 3e, as a chromosome territory example – fit in an averaged 10-micron sized nucleus? A back-of-the-envelope calculation, using average chromosome 10 for example (assume that all 46 have this vol.), with a cylinder representation (1 micron diameter by 10 microns length), gives a summed volume of all 46 chro3osomes of 361 µm^3^. This compares to a 10-micron nucleus diameter volume of 524 µm^3^, leaving one quarter of the nuclear volume free. Thus, this model of a chromosome territory is reasonable.

## Results Part IV: Extensions to the Unified Interphase Chromosome Structure

### A Polytene Chromosome Interphase Chromosome Example: Unified Interphase Chromosome Structure

One of the interesting features of the S chromosome structure is that it is able to unify other interphase chromosome structures. One of these interphase chromosomes, architecture unsolved, is the polytene chromosome organization, seen in many insects and plants (39). These cells/nuclei are locked in G1/S such that the DNA replicates many times, but the cells do not divide, and the chromosomes become large banded (about 200 nm band thickness) chromosome structures (39, see Fig.4).

**Figure 4.**
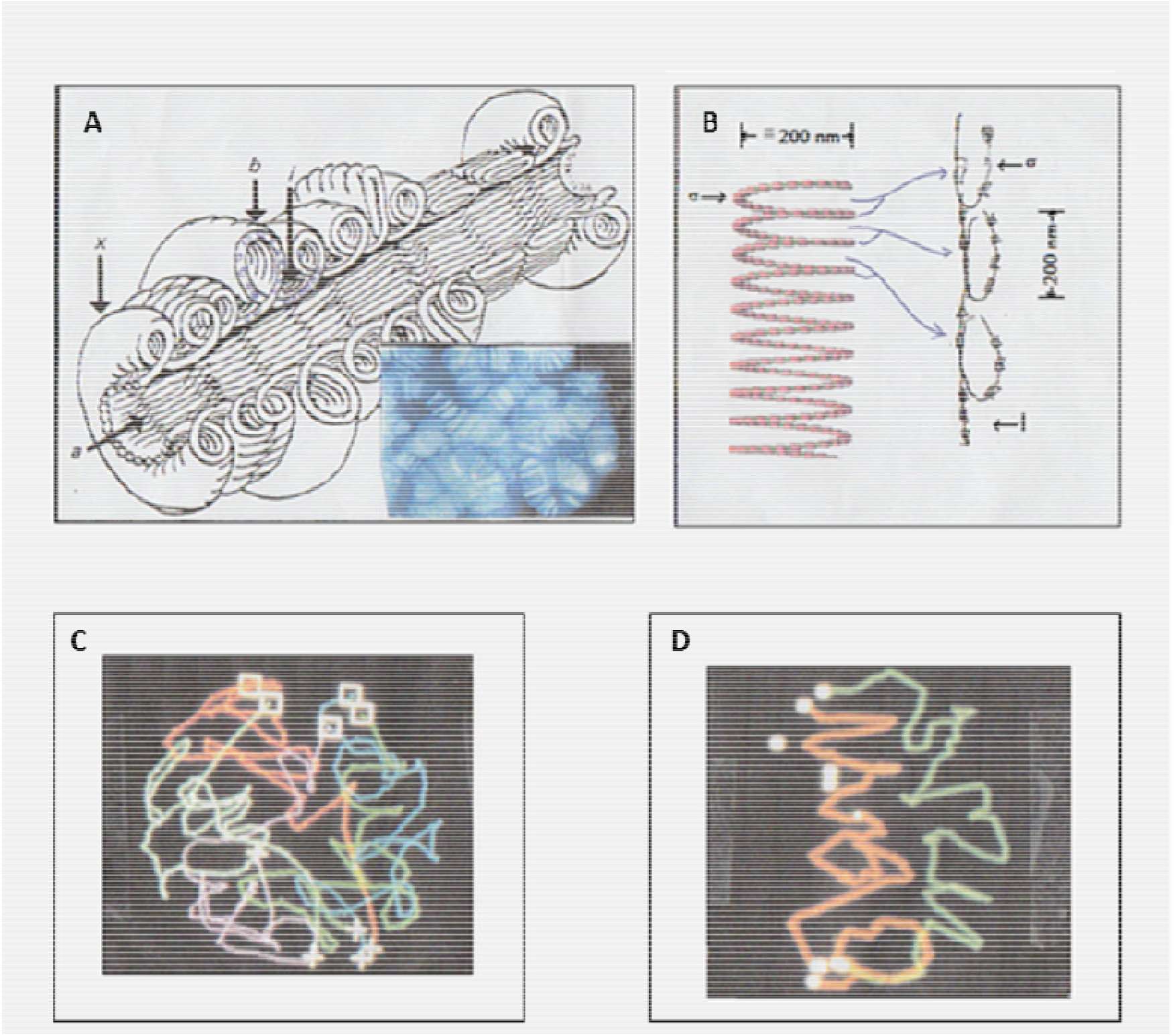
Polytene chromosome architecture can be unified with the diploid interphase chromosome S structure. The polytene chromosome architecture (Fig4a) is taken from Fig 18 of J. Cell Sci. 57, 73-113 (1982), and is reproduced/adapted with permission, as described in the text for polytene chromosomes. Polytenes in the nucleus, as an insert come from the sedat archive. Fig4b shows that each polytene band is possibly a S gyrus (approx..200nm dia) twisted 90degrees as shown giving the approximate dimensions of polytene bands. The polytene banded chromosomes, in a whole nucleus, can be determined, as shown in the reproduced/adapted Fig3 from J. Cell Sci. Suppl. 1, 223-234 (1984) with permission, and are in a Rable configuration (Fig4c). Two adjacent polytene chromosomes from a 3-dimensional nucleus, never mixing with each other, are seen as right handed sloppy coiled structures (41), again in Rable configurations (Fig4d). Data taken from Sedat archive.

**Fig. 5.**
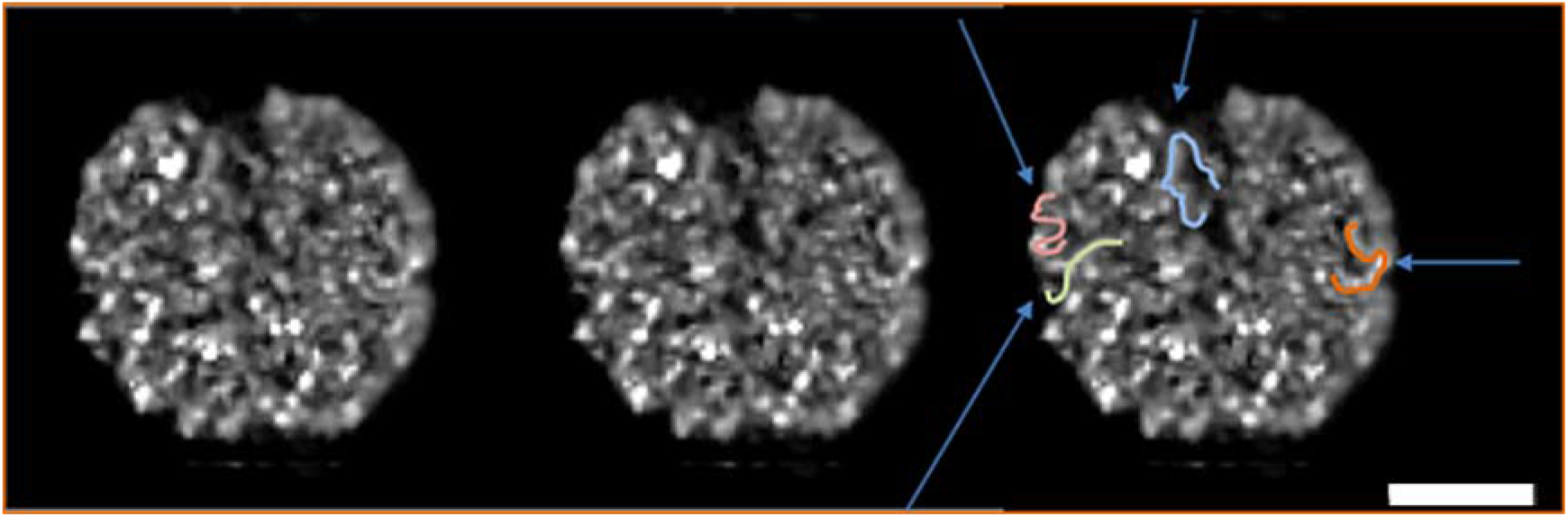
Live optical imaging at extremely low excitation light intensity and DC show unique interphase chromosome structures. The first two images are a stereo pair of a 3-dimensional G1 cell nucleus, visualized by cross-eyed stereo technology (4,5). The third image is a duplication of one of the stereo pair, but used to show, by blue arrows, locations of some of the large scale looped structures, and drawn chromosome looped structures as examples. Scale bar is 4 microns.

Fig. 4A shows a proposed polytene structure, extensively documented, and described in a 1982, in the last century, paper (40). The 1024 DNA copies are confined to a chromosome rim, a torus-radially symmetric chromosome architecture, with a hollow interior. EM studies showed that the path of the DNA, as a fiber undefined then, is likely a single loop about 200 nm side, the band thickness, and diagrammatically shown in Fig. 4A (and 40). It is inferred that the fiber is likely today a NS 11 nm fiber. We propose that the single loop is a S gyrus, same size, rotated 90 degrees as shown in Fig. 4B. The polytene band thickness is the same as a S turn, as shown. Once the loop is rotated, it replicates radially with the same size, and the polytene band growing in width (3 microns for *Drosophila* salivary gland polytenes) as replication rounds proceed. For completeness, Fig. 4C shows a polytene 3-dimensionally determined (41), with Fig. 4D two side-by-side polytene chromosomes, right-handed helically coiled, in the Rabl configuration (37). Thus, it is possible to unify – here-to-fore unsolved – diverse interphase chromosome architectures into a common S molecularly identical structure.

### Lampbrush Chromosome Structure Can Be Integrated Into the S Architecture

Lampbrush chromosome structure has always been difficult to integrate into general interphase chromosome architecture. This enigmatic diplotene meiotic prophase I stage, with paired half-bivalents (2 sister chromatids), has extensive transcriptionally active loops bounded by dense chromatin regions (42, 43 and references therein). Nevertheless, this structure can be readily integrated into the S architecture. We propose that the large transcriptionally active loops are just looped-out single (or several) S helical gyri, where the loop bases are confined, adjacent to more condensed S helical gyri (like the proposed more silent heterochromatin S regions). The transcriptional looping is a natural feature to the S architecture as transcription first extended the S gyri, further with more transcription, with the extension limit being a large loop. Thus, S architecture unifies again interphase chromosome structure.

### A Confirming Living Light Microscope Cell Study

The last piece of evidence for a specific interphase nuclear chromosome structure comes from live imaging of a B cell in likely G1 cell-cycle state (30). The dominant problem in live imaging is phototoxicity from the photons used to excite specific fluorescence giving rise to free-radicals (31, 32), and the repair of the photo-damage that complicate the interpretation of the resulting image. To avoid photo-damage, the conventional excitation intensity is reduced by three to four logs – approximately 1/1,000 to 1/10,000 the usual intensity/dose of photons. The resulting images, taken as a 4-D data stack are extremely noisy (see methods), and computer processing with the newly developed entropy regularized deconvolution algorithm (33) recovers the image as shown in Fig. 5, as described in the methods. The interphase nucleus is fluorescently labeled, so the chromosomes are visible (see discussion in Methods), but best studied as a stereo pair (Fig. 5 and its legend). Study of the chromosome structure in the nucleus suggests that the entire nucleus is filled with a relatively uniform thick looping rope-like structure whose diameter is approximately 260 nm (measured from the size bar). The thick fiber – pointed out by arrows in many places – has small variations or protuberances – bulges – along their entire length. The fiber makes rapid bends and twists, and is packed fairly densely, making close contact to other fibers, approximately one or two fiber diameters away on average. The fibers touch one another in many places.

In summary, evidence from living cells, minimally perturbed, suggests that a G1 interphase chromosome structure is a thick relatively uniform diameter fiber, and will be shown below to structurally match the nuclear cryo-EM chromosome structures. The live nucleus study presented here is suggestive, but the majority of the evidence for a interphase chromosome structure will come from the CET study detailed above.

## Discussion

We have made the case for the existence of a unified interphase chromosome structure, the Slinky, throughout the interphase nucleus. Key to the analysis leading to this structure was the visualization techniques of the CET DC data. A few points emphasizing issues and features can be made: First, this is a defined structure, an architecture using coiling as structural principle. The S structure, with its possible intrinsic order (see Fig. 3b) – really analogous to a molecular antenna – lends itself for further protein–protein interactions almost like a crystalline underlining architecture for further modifications. To put in perspective, no papers have found evidence for the 30-nm NS structure (22, 25) – an in vitro structure – in vivo, in living cells (23, 24). No statement as to coil handedness is made, though this feature could be differentially used.

How are the S structures built or maintained? While no statement can be made; though the polytene literature suggests that each polytene chromosome band may have specific protein(s) to individualize these structures (39), and perhaps this characteristic is in common to interphase S. Cohesin complex with ability, by its ring structure, to bring two structures together (44–46), allows binding, and to bring two adjacent S gyri together, thus further organizing the S. The condensins could change the S architecture (47, 48). Nevertheless, there is a great deal of flexibility and room for genetic modification; the S gyri do have substantial size variation, which could have genetic ramifications, for example for genetic control/distinction. We are reminded that polytene chromosome bands have distinct thickness, thinness, size, and appearance variations – the genetic location band maps – all likely under genetic control (39). Recent literature, the beautiful *Drosophila* sequence and genetic link-up papers, strongly suggest that the polytene band organization – bands, specific bands, and interbands – are highly correlated with the molecular and sequence information (49–51). We make no statement where promoters, enhancers, or protein coding sequences are located relative to the S gyri, though these features could be coupled into the S organization for further genetic specificity.

Second, there is a great deal of variation in appearance of the S bands; bands are bent, twisted, and misshapen, not surprising due to the solution and motion forces that take place in the nucleus. Metabolism issues, transcription and/or replication with their plethora of protein complexes must change significantly the S structure, and this is what we see in the experimental data. Thirdly, the S structure is very flexible and easily bent and moved around, but is this a resulting polymer solution following polymer statistics? Or are the S themselves moved around to specified locations/long range interactions. Indeed, the packing density for the LSS, looking at the data, is high, though one sees LSSs on top of each other in cases, possibly reflecting the possible familiar TADs (28, 29). Tangles, knots, or complicated looping configurations are not seen, though discussed in the literature (52, 53). The LSS are seen to be smoothly aligning and associating with one another. Fourth, the S structure is a delicate structure, easily distorted and modified by chemical fixation, raising issues with the conclusions for these procedures. Fifth, while there is a preponderance of evidence, both live optical data and the CET studies, leading to a defined chromosome structure, there is a great deal of unknowns allowing additional structure(s) to be superimposed. Additional variants might be seen in the different parts of the cell cycle. The poor definition of the cell cycle point for the data presented here may introduce some questions for structure variations.

Lastly, why do we believe this structure out of many in the literature (54, 55)? Simply, we emphasize that the cryo-preservation, in a vitrified aqueous environment, preserves the likely living structural state.

In summary, preliminary evidence, from a CET nucleus data set, was presented showing that the whole nucleus seems to be organized into a defined, approximately 200 nm diameter, coiled NS interphase chromosome architecture, a structure similar to a Slinky. This structure also matched the live optical microscopy study showing very similar image features and nuclear chromosome packing, as seen in the results. This flexible structure allowed a diverse interphase chromosome state, the polytene chromosome state or lampbrush chromosomes, to be unified and understood as essentially the same organization. Thus, all interphase chromosomes might have essentially an identical molecular underlining architecture.

## Materials and Methods

### Living nucleus imaging study (Fig 5.)

Cells were prepared as previously described (30). Briefly, Pro-B cells were purified from the bone marrow using B220 microbeads (Miltenyi Biotech) from mice that carry a tandem repeat of TET repressor elements in both alleles of the immunoglobulin heavy chain locus. Isolated pro-B cells were grown for five days in the presence of interleukin-7 and SCF at 37°C under 5% CO_2_ conditions. Actively growing pro-B cells were transduced with virus expressing TetR-EGFP. Two days post-transduction cells were imaged using the OMX (Applied Precision, Issaquah, WA 98027, V3 (3.20.3537.0), vintage 2010) microscope platform. Typically, cells could be distinguished in G1/perhaps early S from later cell cycle stages by whether there were one or two Tet dots. Images were acquired using a 100x/NA 1.4 objective using 100 time points (each time point for 10 ms, 1 s/data cube), 0.15 micron Z step (typically 30–50 Z sections/cube), 2 OD/0.01 intensity attenuation, and a 5 ms exposure time. Even with the EMCCD (with high gain) as camera image acquisition, the individual Z sections, before DC, showed just noise; Z projections of the data could faintly just see that a nucleus was present.

There are some qualifications to the live nucleus chromosome study. First, we note that the G1 cell-cycle state is approximate. Second, the fluorescent chromosome label comes from low-affinity interactions involving TET-GFP repressor binding across the genome. Third, it will be important in future to examine TET-GFP mediated fluorescence across chromosomes using a range of intensities. Finally, we emphasize that this live chromosome data set is merely a supporting example to the CET data that is the majority of evidence for the S structure proposed in this publication.

DC of the Fig. 5 light micrographs, as 3- or 4-dimensional images, were done under ERDecon-II (33, extended/updated by MA and Eric Brandlund), now in Priism (see below). DC followed (33), but in some cases an additional time (second derivative) filter was used; in ERDecon-II the time filter was a natural extension to the DC software, built into the mathematics of ERDecon-II (33). The improvement of the 4-dimensional over the 3-dimensional DC was slight, and Fig. 5 is the 4D version.

### Initial DC Processing

The data for this paper comes from an earlier publication (3). The cells were grown, plunged frozen, and processed for CET as described in detail in (3). The tomograms were DC as described in (3), and an initial representative nuclear DC CET image is Fig. 6 in ref. 3.

This nucleus CET was further processed and analyzed at UCSF. First, the DC data was intensity scaled to remove(truncate) the pathological, sparse, isolated, few, extremely high intensity pixels from the DC data (see supporting Information section). Next, the DC image was histogram trimmed (HT), a process specific to DC images. The DC intensity histogram typically has a lower intensity values (mostly noise and poorly DC), while the upper intensity values, the broad higher intensity tail, has all (most) of the DC image/structure features (much better DC – largely filling the missing wedges that were missing – (see 3,60). There were, in addition, also quite large intensity values coming from the calcium phosphate grandules in the mitrochrondria. The low and high intensity values were then truncated. This process is discussed in detail, with images, in the Supporting Information section.

The HT DC image was intensity displayed, in 3 dimensions, using Quick-Time display software present in every computer (see Fig. 1). Z planes can be studied (by moving up and down Z in the cube), selected by movement of the cursor bar. The majority of the data for the interphase chromosome structure comes from the stereo rocking angular movies (RASPs), described in detail in the Supplement (initial examples are seen in 60). The RASPs allow 3-dimensional views of dense cellular structures for detailed study and analysis. The computer scrips (written in Priism–see below) are available from the authors(JS and AM).

### Computer Modeling Software

The computer modeling of the MC, at all levels, utilized a software package written by a Turkish Engineering group (57) that allows sequential helical coiling of defined sized structures. This package is run under, and requires, Mathematica12.3 (58) under Linux in large work stations; Mathematica12.3 was run under Windows 7, Intel Core i5 CPU 3.20GHz processor with 4 cores and displayed on a Samsung C27F591 monitor and NVIDIA Quadro K1200 video card and 8GB of memory. Mathematica was also run on a CentOS 7.3 Linux system running on a Intel Core i7 CPU 2.80GHz processor with 16GB of memory. The display of the Linux generated output was visualized on the Windows 7 computer. The Turkish software (57) was modified to take into account the RT S’ configuration modifications. The various scripts for the software are supplied by the authors upon request.

### Computer Display Software

Once structures are built, they are displayed, in various dimensions, with the generalized display and quantitation software package written over the years by the Agard/Sedat groups (59). This software, and its extensive Help files, is available from the Agard and Sedat UCSF emails. The display scrips are also available from the authors.

## Supporting information

RASPA

RASPb

RASPb'

RASPb''

RASPb'''

RASPc

RASPc'

## ACKNOWLEDGMENTS

OMX use was made possible UCSD school of medicine Microscopy Core Grant P30 NS047101. We gratefully acknowledge the advice and suggestions of Professors Markus Noll, Lloyd Smith, David DeRosier, Robert Stroud, Marc Shuman, David Agard, and Eric Brandlund. Research in the Murre laboratory is supported by the National Institutes of Health (AI082850, ROAI00880, and AI09599). ME is incumbent of the Sam and Ayala Zacks Professorial Chair in Chemistry. His work is supported by a grant from the Israel Science Foundation (1696/18). Fig 2. Used UCSF Chimera, developed by the Resource for Biocomputing, Visulization, and Informatics at the University of California, Sam Framcisco, with support from NIH P41-GM103311.

