## Supplementary text file for "A Proposed Unified Interphase Nucleus Chromosome Structure: Preliminary Preponderance of Evidence"

**Supporting Information**

**Essential movie data**

The movie data, in files, will need to be downloaded to the desk-top for ease of access.

First, there is Fig. 1, a Quick Time 3-dimentional data stack that is clicked on and displayed in Quick Time, present in all computers, with a bar, that can be displaced, to move up and down the Z axis.

Second, there are the 9 stereo movies, the Rocking Angular Stereo Pair (RASP) files [RASPa–c] which are played by clicking the file name and looking in stereo as detailed below. These are essential to study the 3-dimentional data that is the basis of the evidence for the interphase chromosome structure.

**Special Nuclear Structure Visualization Problems and Their Solutions**

One of the best ways to study and analyze the nuclear CET chromosome structure is through the use of stereo pairs (SP) (1,2) to take into account the intrinsic 3-dimensional nature of the data. SPs are best seen as side-by-side images fused after crossing one’s eyes. A tutorial below will outline how to rapidly see stereo by the crossed eyes technique. A second tutorial is provided to further aid in visualizing the 3D interphase chromosome structures, in the nucleus, once the stereo fused images technique is learned from the first tutorial.

Usually, a small cube of data is computationally cut out of the DC for SP study. The SPs are further modified by rocking the SP so that the 3-dimensional data can be seen from different angles, typically 180 degrees of tilt. In addition to rocking, the SP data cube is rotated by large different (Euler) angles to further displace the 3-dimensional structures. These SP rocking and large angle views are called RASP (defined above), and will be one of the primary ways the chromosome structure will be analyzed.

Further SP modifications may be necessary for structure analysis. Viewing the SP movies can, in many cases, be enhanced by variation of the size (can be modified by movement of the lower right-side diagonal tab), so that structures can be discerned, differentially, if viewed at a distance which size allows one to do. Motion, rapid rocking of the views, can in addition be used to discern additional structural features (the structure can snap into view with motion). The RASP movies, because both eyes see features slightly differently, give the impression of enhanced (better) resolution. One usually requires that the same structure is seen at close adjacent (fast) rocking angles, and in many cases that structure can be reconciled/seen at the other large angular views to be believed. The RASPs turn out to be crucial for the interpretation of the banded LSSs as will be demonstrated. It is worth pointing out that there is a learning curve for the study of the RASPs. Video movie 2 of the Supporting Information is a guide to seeing the structures in the nucleus RASP movies. Initially, five to ten minutes of study are usually required to see the banded structures reliably, but subsequently they are rapidly found in abundant places.

Finally, RASP(s) are studied for each nuclear box in Fig. 1. The detailed LSS banding and its sub-structure is really only appreciated by these SP movies. In some cases these nuclear boxes are shifted in Z, giving rise to slightly different RASP(s), for example Fig. 2b B’–B’’’, so that the banded LSS at different Z depths (and their modifications) can be studied.

**Problems for LSS Data Interpretation**

Before interpreting the LSS and their bands (presented in the Results section) one needs to discuss a number of general interpretation problems. First, EM grey-level data, both for STEM and TEM, represents mass; though, STEM via scattering is rigorous mass (3). However, the grey levels lack specificity, and one cannot say which grey level is which, protein or nucleic acid or their complexes. Instead one uses molecular attributes and organelle locations to define structures, a crude process (4). There are ways to think about this problem, and the solution is in the future (5,6).

While the chromatin/nucleosome makes up the bulk of the nuclear mass, it is not possible to rigorously specify these structures. Likewise, local densities, slight thickenings, etc. could represent transcription products, transcription factor complexes (usually multi-protein complexes), transcription products, processing/splicing factor complexes, or other structural components. A second problem is that one typically needs to scale the images for intensity, hence mass. The grey levels are in a very large dynamic range that cannot be all displayed. Instead, upper and lower intensity levels are displayed, but in the process some structure/mass is clipped out, thereby missing possible structural features. In several cases, several dynamic ranges are simultaneously studied to remedy some of these problems. Coupled with this problem is the worry that very weak features – such as the 2-nm bare DNA fibers – are not visible because of the scaling; in addition, one worries that structural features are lost because one is at the end of the CET resolution.

A third problem is geometry considerations for the LSS. It is a flexible, bending, twisted, dotted- (broken up because of sub-structure), and a possibly coiled structure. When it is studied by selecting Z planes or sub-regions, these structures are computationally cut so that different orientations are seen in complex geometries, complicating the analysis. In addition, these structures can be on top of one another in some places in the nucleus. Fourth, there is the problem of where one is in the cell cycle. One does not know if the nucleus is in G1, S, or a mixture, complicating the interpretation of the structures. This issue will be resolved in future CET studies.

**Stereo pair viewing tutorial**

A training stereo pair is shown in Fig. S1. This image is downloaded as a side-by-side horizontal pair about 4–5 inches across separated by 1–2 inches. Next this pair of images are studied/viewed approximately 12–18 inches away (straight above the images), crossing one's eyes until a third picture between the two pictures appears, then gently relaxing ones eyes, but still slightly crossing them, so that the third image appears in the middle between the two pairs. One concentrates on this third image, and immediately a stereo 3-dimensional, dominant image appears. In the trial Fig. S1, these geometric structures in 3-dimensions are evident, spaced 3-dimensionally. Then one continues to train by seeing stereo rapidly, then experimenting by viewing the stereo images increasing distances away ( 24+ inches away), and even viewing off axis. Large-scale 3-dimensional structures can be discerned, in some cases, by viewing 6–8 feet away from the stereo pairs. See <https://en.wikipedia.org/wiki/Stereoscopy> for general stereo viewing.

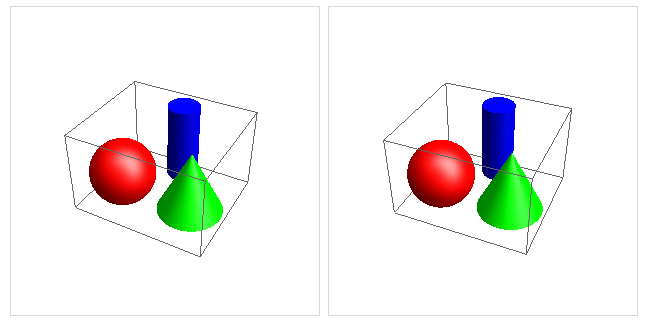

Fig. S1. Stereo training geometric models

3D geometric figures [<https://mathematica.stackexchange.com/a/30015/77273>]

A second training image is provided as a firework movie [<http://youtu.be/dpyjYWfZESw>] so that time dependent 3-dimensional data are seen in stereo as above.

**Stereo viewing tutorial for RASP movies**

Now that it is possible to view stereo in a facile way, we can proceed to the RASP data movies. Start with RASPa and be able to see the different angular views that are rocking by the cursor movement, and be able to jump from one view to another in stereo, seamlessly. Be able to reconcile a structure from one rocking and a different angular view. This ability may take a few tens of minutes to accomplish.

Next study features, like heterochromatin, in Fig. 1 and Fig. 1-guide to pick out 3-dimensional chromosome structures. Look for repetitive structures that are perpendicular to the long axis of the LSS, that are curving. Next look for sub-structural features like bumps and sub-structure with grey level differences/texture in 3 dimensions. Lastly, study the features in Table 2 using these new visualization techniques. Many additional examples will be discerned.

**DC processing; Histogram Trimming Details**

The DC post processing and the HT is now described in detail. First, the DC 3-Dimensional image from (7) had the pathological intensities (~500,000) removed; the intensity values from 8000 and above were truncated to 8000 (Fig S2B.). Second, the low intensity values,less than 1500 and the values greated that 8,000 ( the high intensity values come largely from the mitrochrondria granules), part of the HT process, were truncated to their set values ( 1500 & 8,000 ), as shown in Fig S2C.). This HT process is described in detail in (7) Fig. 5 and Fig. S4 in the Supplement for (7). Finally, this DC image was trimmed in X,Y , and then Z was trimmed above and below the nucleus to center the nucleus as shown in Fig S2D.

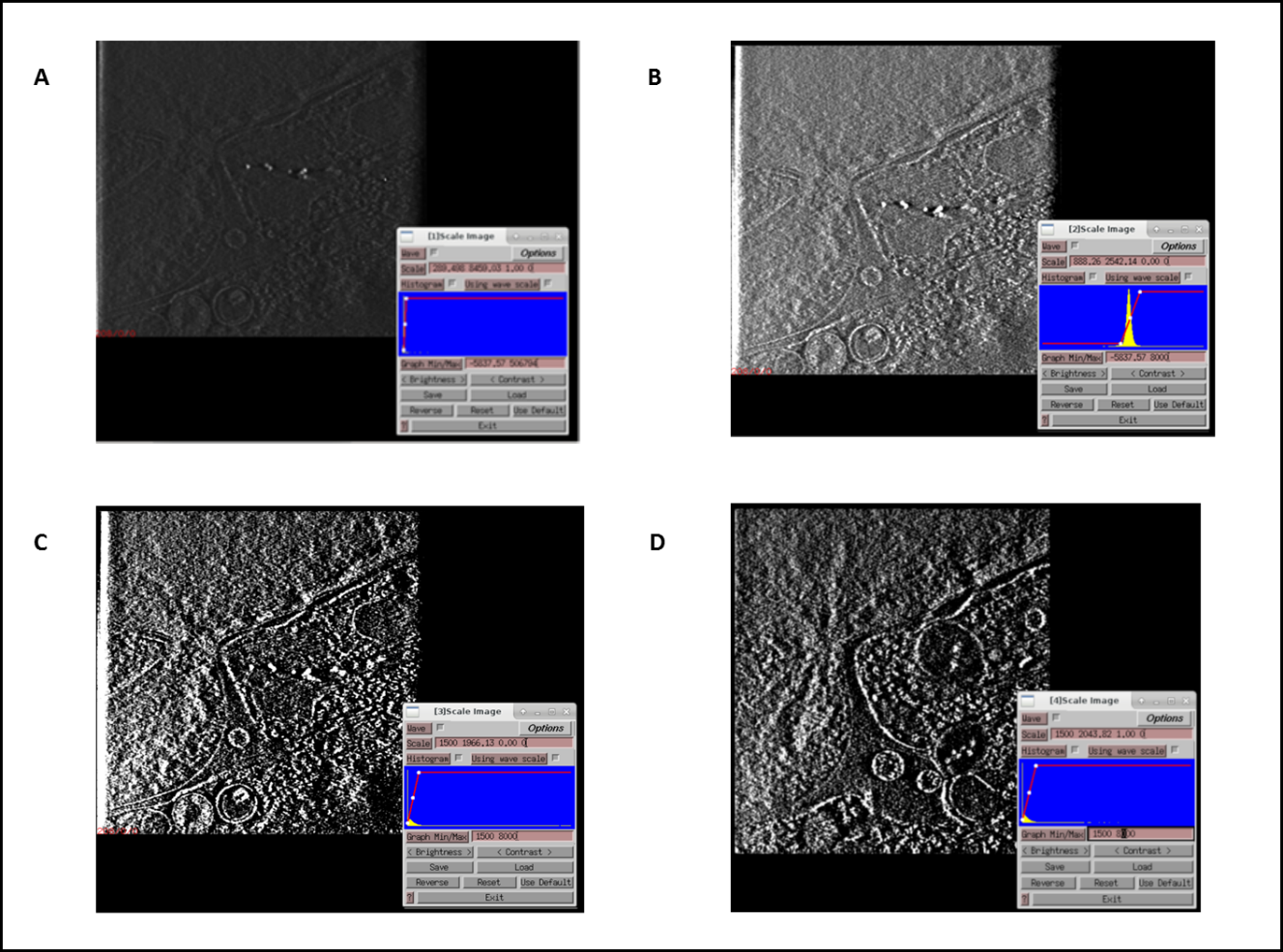

**Fig S2**. Different DC HT and post-processing steps

Fig S2A shows the DC image as it came from (7) with the intensity range of ~(-5000) to ~500,000 as shown in the histogram insert. Fig S2B is A with the intensity values greater than 8,000 truncated to 8,000. FigS2C is B with the lower than 1500 intensity values truncated to 1500; the intensity range is now 1500 to 8000. Fig S2D has the XYZ borders trimmed ( pixels removed) to center, and only included nucleus DC structures. The intensity values go from 1500 to 8000.
